# Lymph Nodes Inhibit T-cell Effector Functions Locally by Establishing Acidic Niches

**DOI:** 10.1101/689604

**Authors:** Hao Wu, Veronica Estrella, Pedro Enriquez-Navas, Asmaa El-Kenawi, Shonagh Russell, Dominique Abrahams, Arig Ibrahim-Hashim, Dario Longo, Yana Reshetnyak, Kimberly Luddy, Mehdi Damaghi, Smitha Ravindranadhan Pillai, Matthew Beatty, Shari Pilon-Thomas, Pawel Swietach, Robert J. Gillies

**Affiliations:** Dept. Cancer Physiology, H. Lee Moffitt Cancer Center and Research Institute, Tampa, FL 33612, USA; Cancer Institute, The Second Affiliated Hospital, Zhejiang University School of Medicine, Hangzhou 310058, P.R. China; Dept. Cancer Immunology, H. Lee Moffitt Cancer Center and Research Institute, Tampa, FL 33612, USA; Dept. of Molecular Biotechnology and Health Sciences, Univ. of Turin, −10124 Turin, Italy; Dept. Physics, Univ. of Rhode Island, Kingston, RI 02811, USA; Dept. Physiology, Anatomy and Genetics, University of Oxford, Parks Road, OX1 3PT, England

## Abstract

Lymph nodes are an essential component of the adaptive immune response where antigen-presenting cells are closely housed with their cognate effector cells. Protection of lymph node resident cells from activated immune cells in such close quarters would need to be robust and reversible. Effector functions of T-cells are profoundly and reversibly inhibited by an acidic microenvironment. The underlying mechanisms of this inhibition are unknown, but may relate to glycolysis, which is obligatory for expression of effector functions. Here, we demonstrate that acidification rapidly and potently inhibits monocarboxylate transporter-dependent lactic acid efflux, which dually inhibits glycolysis by end-product accumulation and by reducing cytoplasmic pH. Based on the robustness of these responses, we propose that acid-evoked T-cell inhibition is physiologically important, and that lymph nodes are a natural site for such modulation. Using multiple imaging techniques, we show that paracortical T-zones of lymph nodes are highly acidic. We further show that T-cells can be activated by dendritic cells at low pH, and their effector functions are restored rapidly upon increasing pH. Thus, we describe a novel physiological mechanism whereby activated T-cells are kept in stasis by acidosis whilst resident in lymph nodes.

## Introduction

It is known that CD8+ effector T-cell functions are inhibited by low pH(*1–3*). For example, CD3/CD28-activated T-cells from C57BL/6 (B6) mice had dramatically reduced interferon-gamma (IFNγ) release when incubated for 24 hours at pHe 6.6, compared to time-matched cells incubated at pHe 7.4 (**Fig. 1A**). In a large panel of cytokines, secretion was generally reduced by acidity, but this effect was not due to a generalized failure of exocytosis because the release of some cytokines (MDC, MIG and IP-10) was increased at pHe 6.6 (**Fig. 1B; Fig. S1A**). Attenuated secretion of IFNγ was also not due to acid-catalysed conformational disruption, because IFNγ immunoreactivity remained stable over a wide range of pHe (**Fig. S1B**). Acid-inhibition of IFNγ and interleukin-2 (IL-2) elaboration was titratable in T-cells isolated from four different strains: *(i)* B6 T-cells stimulated with anti-CD3/anti-CD28, *(ii)* OT-I (CD8+) T-cells stimulated with OVA_257-264_ SIINAFEKL peptide, *(iii)* PMel (CD8+) T-cells stimulated with gp100 peptide, and *(iv)* OT-II (CD4+) T-cells stimulated with OVA_323-339_ ISQAVHAAHAEINEAGR peptide (**Fig. 1C/D**). Previously, we reported a build-up of IFNγ mRNA in stimulated PMel T-cells under acidic conditions(*2*), and we postulated that this would “prime” cells to express effector functions, when returned to alkaline conditions. Although we have confirmed this observation in PMel cells, the effect of pH on IFNγ mRNA was not consistent among the other types of T-cells (**Fig. 1E**). If T-cells are stimulated for 24 hr at pH 7.4, and then incubated at pH 6.6 for 24 or 48 hours, they rapidly (blue lines) release IFNγ production following re-stimulation at 7.4 (**Fig. 1F**), suggesting that, despite that effects on mRNA, priming may indeed occur.

**Figure 1:**
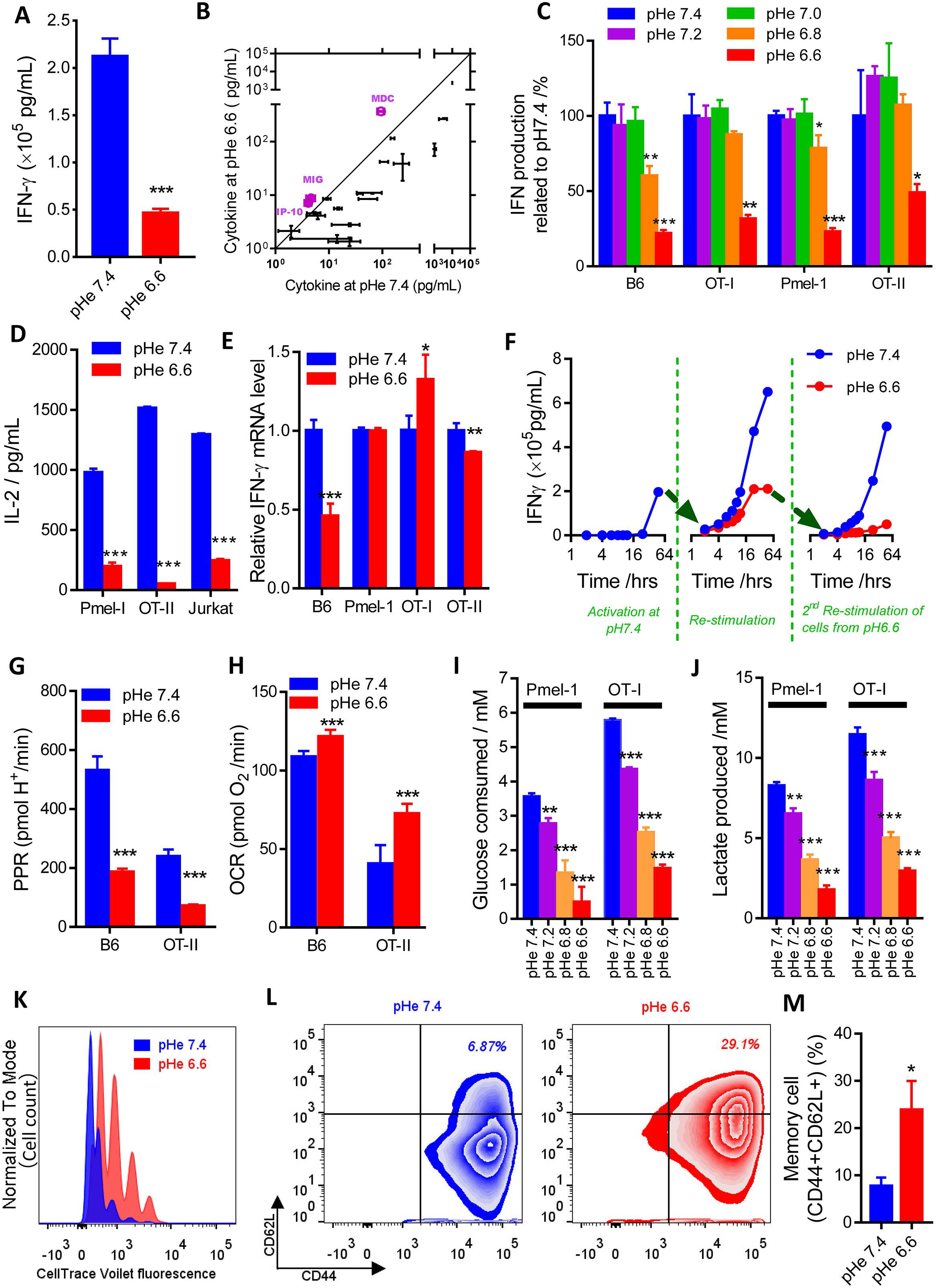
T-cells effector functions are inhibited at acidic pH. (A) Reduction in interferon γ (INFγ) release from B6 T-cells at low pH measured by ELISA. (B) Relationship between cytokine levels determined in paired experiments at low and high pH, measured by Cytokine Beads Array (CBA) assay, showing reduced release in acidic conditions for most cytokines, with the exception of those shown in purple. (C) INFγ production over a range of pHe, normalized to levels probed at pHe 7.4 in four types of T-cell. (D) Interleukin-2 (IL-2) release (ELISA) from three types of T-cell or leukaemia cell line is reduced at low pHe. (E) The effect of low pHe on INFγ mRNA measured in four types of T-cell, showing cell-dependent behaviour. (F) Time course of INFγ levels in media following a pH-manoeuvre. Acid-inhibition of cytokine production can be reversed upon exposure to alkaline pH. (G) Proton production rate (PPR) is reduced under acidic conditions. (H) Oxygen consumption rate (OCR) is increased under acidic conditions. (I) Glucose consumption and (J) lactate production as a function of pH. (K) Rate of B6 cell proliferation measured by CellTrace Violet is reduced at low pH. (L) Representative data showing shift towards memory phenotype of T cells at low pH. (M) Fraction of CD44+CD62L+ cells (i.e. memory phenotype) is increased at low pHe. Mean±SEM of 5 experiments. All the experiments were repeated at least twice and expressed as mean±SD (unless indicated otherwise). Significance level: *, p<0.05; **, p<0.01; ***, p<0.001.

T-cell activation evokes a dramatic increase in aerobic glycolysis, which is necessary for enabling effector T-cell functions(*4*). The underlying mechanism involves the glycolytic enzyme glyceraldehyde 3-phosphate dehydrogenase (GAPDH), which is diverted to bind to IFNγ mRNA and suppress translation(*5*) if glycolytic flux is low. Glycolytic flux can be quantified using a Seahorse extracellular flux (XF) analyser, which measures an extracellular acidification rate (ECAR) that can be converted to a quantitative proton production rate (PPR). At 48 hrs following T-cell activation at pHe 7.4, the PPR of B6 and OT-II T cells were 240 and 530 pmol H^+^/min (ca, 5-15 mmol/L cell) and these were severely attenuated at pH 6.6 (**Fig. 1G**). The basal oxygen consumption rate in the presence of glucose was incrementally increased (**Fig 1H**), consistent with a depolarization of mitochondrial membrane potential (**Fig. S1C-D**). Glycolytic inhibition at low pHe was confirmed through measurements of glucose consumption and lactate production in media (**Fig. 1I, J**). Inhibiting glycolysis following T-cell receptor (TCR) engagement, has been reported to promote differentiation into memory T-cell phenotypes(*6*). In our system, acidosis decreased the proliferation rate of T-cells (**Fig. 1K**), including both CD4+ and CD8+ subsets (**Fig. S1E**), and promoted the differentiation from an ‘effector’ to a ‘memory’ phenotype(*7*) (**Fig 1L,M**). The later observation is consistent with the inhibitory effect of acidity on mTORC1 activity(*8*) that can induce memory T-cell differentiation(*9*).

The mechanism through which acidity influences T-cells is not known, but many H^+^ ion effects involve reversible binding to histidine residues in proteins(*10*). Thus, there is a myriad of possible targets, including, *inter alia,* activation of acid sensing receptors or channels(*11*), modulation of Ca^2+^ signalling(*12*), and changes in intracellular pH (pHi) affecting intracellular targets(*13*). Two acid-sensing G-protein coupled receptors, GPR65 (TDAG-8) and GPR68 (OGR-1), are expressed in T-cells(*2*), but activated T-cells obtained from *Gpr65* or *Gpr68* knockout mice remained inhibited by acidosis (**Fig. S2A**). Furthermore, small-molecule inhibitors of OGR1 and GPR4 (BA-39-PQ30-1, NE-52-QQ57-1, gifts from Novartis) failed to rescue cytokine release under acidic conditions (**Fig S2B-C**). Inhibition of TRPV1, an acid-sensing ion channel, did not rescue IFNγ production, either (**Fig S2D**). Acid-sensing ion channel (ASIC)(*14, 15*) isoforms 1 and 3 are also expressed in T-cells(*2*), but the potent and specific inhibitors, A-317567, APETx2, and psalmotoxin(*16*),(*17*) were unable to restore T-cell function at low pHe (**Fig. S2E-G**). Similarly, amiloride and cariporide showed no ‘rescue’ effect (**Fig S2H-I**). The only treatments that modestly raised IFNγ production at acidic pH were phorbol esters and histone deacetylase inhibitors (**Fig S2K, L**). T-cell Ca^2+^ signalling may respond to low pHe through the pH-sensitivity of Ora1 Ca^2+^ channels. However, low pHe did not meaningfully change store-operated Ca^2+^ entry interrogated by a standard protocol (**Fig. S3A-C**).

Given that metabolic changes are necessary for effector T-cell functions, a plausible link with pHe could be the acute inhibition of glycolysis by H^+^ ions. This was tested by measuring the effect of a prompt pHe change on the ECAR. Injecting a volume of HCl that reduces the pH of lightly HEPES/MES-buffered medium from 7.4 to 6.6 triggered a rapid fall in ECAR in activated OT-1 or B6 cells, which was reversible by injecting a volume of NaOH (**Fig. 2A**). To measure the effects on intracellular pH (pHi), T-cells were loaded with the pH-reporter dye cSNARF1 (calibrated with nigericin, **Fig S3D**) and imaged confocally. Changes in pHe could be produced by switching between HEPES/MES-buffered superfusates titrated to pH 7.4 or pH 6.6, and these evoked dynamic changes in pHi in the same direction but with a short delay and reduced amplitude (**Fig. 2B**). To determine if these pHi shifts were stable, T-cells were equilibrated in either pHe 6.6 or 7.4 and their pHi was measured once they reached a new steady-state by both flow cytometry and microscopy. In prior, work flow(*18*) showed only a small fall in pHi (~0.1 units) when dropping pHe from 7.6 to 6.6, and this was repeated here (**Fig. S2N**). However, this technique cannot resolve nuclear form cytoplasmic pH, plus it must be performed in the absence of physiological CO_2_/HCO_3_^-^ buffer. Hence, HCO_3_^-^-dependent transporters that underpin much of the dynamic pHi-regulation are inactive. Because a majority of the intracellular volume of T cells is taken up by the nucleus, T-cells loaded with cSNARF1 were co-loaded with the nuclear dye Hoechst-33342 and imaged confocally under superfusion with 5% CO_2_/22mM HCO_3_^-^-equilibrated buffer. The Hoechst dye allowed masking of the nuclear signal, resolving the signal from the cytoplasm (**Fig. 2C inset**). Upon decreasing pHe from 7.4 to 6.6, pHi stably decreased by ~0.2 pH units (**Fig. 2C**). Although subtle, such transduction of a pHe change into a sustained pHi signal accesses a myriad of protonatable targets in the cytoplasm, including enzymes in the glycolytic pathway(*19*), such as highly pH-sensitive phosphofructokinase, PFK-1 (**Fig. S3H**).

**Figure 2:**
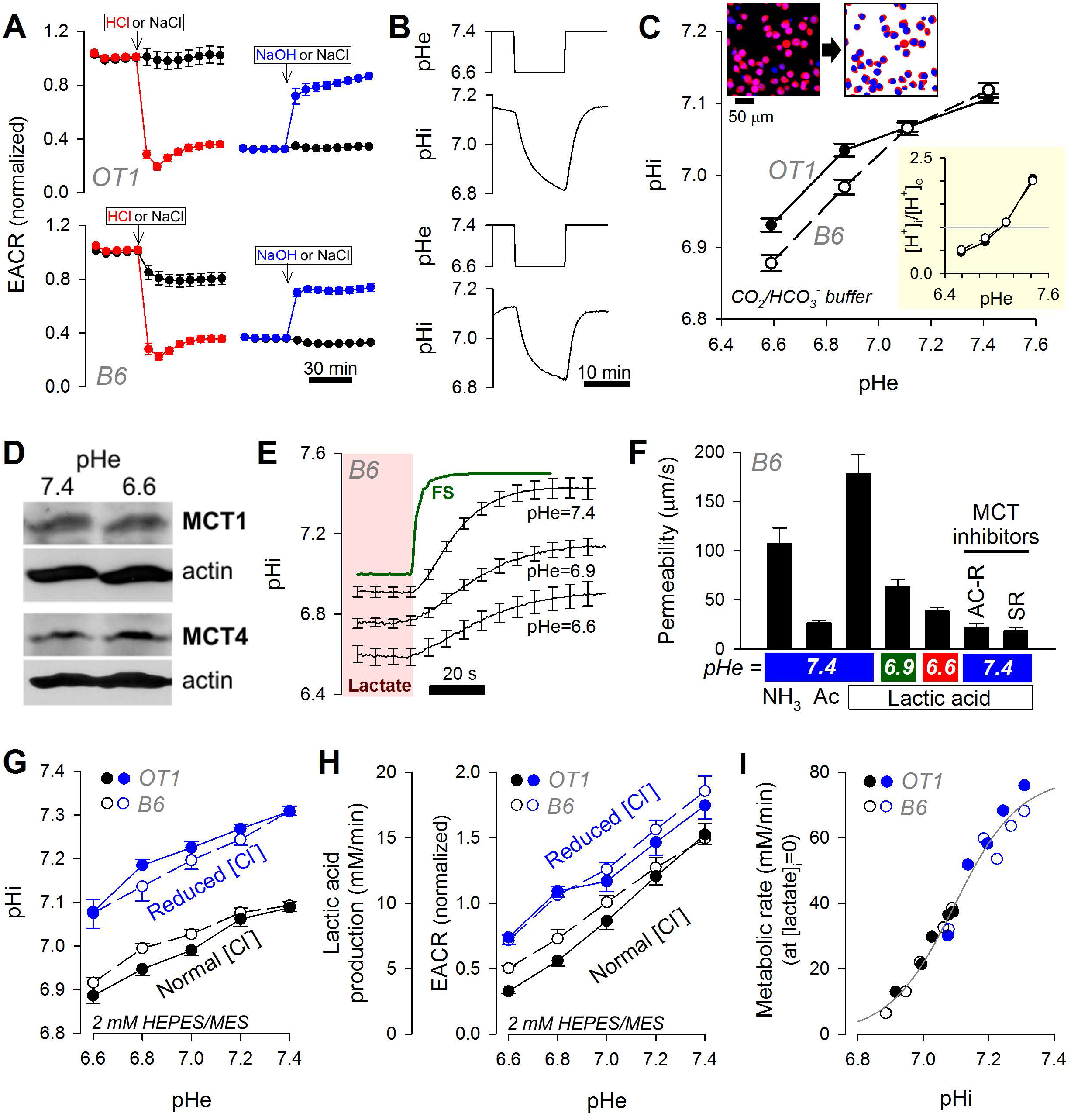
Mechanism of the inhibition of T-cell glycolysis at low pH. (A) An injection of HCl abruptly reduces extracellular acidification rate (EACR) in OT1 and B6 T-cells; this reverses upon an injection of NaOH. NaCl injections performed as sham controls. Solutions were lightly buffered with 2 mM HEPES/MES mixture and titrated to desired pH. Mean±SEM (N=4). (B) A reduction in extracellular pH (pHe) evokes a delayed reduction in intracellular pH (pHi), as measured from cSNARF1 fluorescence under the same buffering regime as used in A. Mean of 10 time course recordings; error bars not shown for clarity. (C) Fluorescence imaging of cells under superfusion with CO_2_/HCO_3_^-^ buffer. Cells co-loaded with cSNARF1 (red) to report pH and Hoechst-33342 (blue) to exclude nuclear areas from the analysis. Plot shows relationship between pHe and pHi at the steady-state in OT1 and B6 cells. Note the transmembrane [H^+^] gradient, shown in inset, inverts near resting pHi. Mean±SEM of 5 recordings of fields of view containing 40-60 cells. (D) Exemplar western blot for MCT1 and MCT4 on lysates collected from B6 T-cells that had been incubated at pHe 7.4 or 6.6. (E) Measuring total MCT activity from the rate of pHi-change driven by transmembrane lactate efflux. T-cells under superfusion were equilibrated with one of the three conditions, 30 mM lactate at pHe 7.4, 15 mM lactate at pHe 6.9 or 7.5 mM lactate at pHe 6.6. Rapid switching to lactate-free solution at the same pHe evoked net lactate efflux. Apparent permeability to lactic acid can be calculated from the rate of pHi change, buffering capacity and transmembrane gradient. To confirm that the ensuing pHi response was not rate-limited by the speed of solution exchange, one solution was labelled with fluorescein sulphonic acid (FS) and the rate of fluorescence-change indicated an exchange time constant of 2.6 s. Mean±SEM of 10 cells per condition. (F) Apparent membrane permeability for NH_3_ (added as 15 mM NH_4_Cl) acetic acid (Ac; 30 mM NaAcetate) and lactic acid (7.5-30 mM NaLactate). Indicated experiments performed in the presence of MCT inhibitors AR-C (AR-C155858; 10 μM) and SR (SR13800; 10 μM). Mean±SEM of 7-15 cells per condition. (G) Steady-state relationship between pHe and pHi mapped for 2 mM HEPES/MES solution containing either normal (140 mM) or reduced [Cl] (7 mM), iso-osmotically substituted with gluconate to offset pHi at constant pHe. Mean±SEM of 6 recordings with 40-60 cells each. (H) EACR, calibrated to units of lactic acid production rate, is not a unique function of pHe. (I) Data from G and H analysed to generate a relationship between metabolic rate, extrapolated to lactate-free conditions (see Eq 1). Best-fit is a simple function of pHi, described by a Hill curve.

Coupling between pHe and pHi arises from changes in transmembrane acid-base fluxes. One such flux is carried by H^+^-moncarboxylate co-transporters (MCTs). Low pHe thermodynamically hinders H^+^-lactate export, leading to an intracellular retention of H^+^ and lactate ions. MCT1 and MCT4 are present in T-cells, and their expression remains stable at low pHe (**Fig 2D**). MCT flux was quantified from the rate of pHi-change evoked by the withdrawal of extracellular lactate, performed using a rapid switching system, which evokes an increase in pHi, the rate of which can be used to calculate a permeability coefficient, P_LAC_ (**Fig. 2E**). The measured PLAC was 180 μm/s in B6 cells (**Fig. 2F**). For comparison, P of acetic acid and NH_3_, which are both freely permeable to the bilayer, were lower (~110 and ~30 μm/s, respectively; **Fig. 2F**), indicating a lactate transport is a protein (MCT) mediated process. MCT1 inhibitors AR-C155858 and SR13800 (10 μM) reduced lactic acid permeability to 20 μm/s, i.e. to the level of protein-unassisted permeability across the bilayer (**Fig. S3F, Fig. 2F**), suggest that it is responsible for P_LAC_. P_LAC_ was also reduced substantially at low pHe (Fig. 2E, 2F), consistent with thermodynamic inhibition of MCT1. Reduced MCT transport kinetics will acidify the cytoplasm and leads to intracellular lactate retention, which feedbacks negatively on glycolysis. Given that the glycolytic flux must be high in activated T-cells(*5, 7*), MCT1 activity is a rate-limiting step that can control T-cell effector functions.

To confirm that a change in pHi is an intermediary in the inhibition of glycolysis by an acidic pHe, ECAR was measured under conditions that could manipulate pHi with and without a concomitant pHe change. Seahorse media were prepared with 2 mM HEPES and MES to provide a low but near-constant buffering. To raise pHi at constant pHe, solution [Cl^-^] was reduced by iso-osmotic replacement with gluconate (**Fig 2G**). This ionic substitution alters the transmembrane Cl^-^ driving force for acid-loaders (e.g. Cl^-^/OH^-^ exchangers) and loads cells with base. Glycolytic rate (J_glyco_) reported as ECAR was not a unique function of pHe (**Fig 2H**); instead, it could be modelled as a simple function of pHi and intracellular [lactate] (**Fig 2I**):

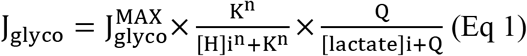

Where K and Q are the apparent binding constants for H^+^ and lactate ions, respectively, and n is the Hill coefficient for the binding of H^+^ ions. Knowing the transmembrane pH gradient (**Fig 2G**), metabolic rate (**Fig 2A**) and lactic acid permeability (**Fig 2F**), it was possible to estimate cytoplasmic [lactate] at steady-state (**Table S1**). By best-fitting these data to Eq. 1, it was possible to describe the relationship between pHi and glycolytic flux in the absence of intracellular lactate, i.e. extract its pHi-dependence (**Fig 2I**). This relationship was found to be highly cooperative (n=4.39) and with half-maximal activation near resting pHi (−log(K)=7.095), i.e. consistent with being a relevant and strong modulatory influence. The ensemble effect of end-product inhibition by lactate was best described by Q of 2.1 mM, i.e. accumulation of lactate to 2.1 mM would halve J_glyco_ at a constant pHi. The model predicts that, at constant pHe and pHi, the addition of lactate would reduce glycolysis by end-product inhibition. Since the L- and D-isoforms are transport substrates for MCT, both will similarly influence transmembrane traffic. In contrast, only L-lactate will produce end-product inhibition on stereo-specific lactate dehydrogenase (LDH). Indeed, the L-isoform produced a stronger inhibition of ECAR to 21% of control (pHi + end-product inhibition) vs. 58% of control (pHi only) with D-lactate (**Fig. S3G**). Consistent with this, Eq. 1 predicts that, at constant pHi, these concentrations of L- and D-lactate would reduce J_glyco_ to 24% and 44% of control.

Since the effect of acidity on T-cell function is so profound, we hypothesised that it has a role in the physiological regulation of adaptive immunity. A plausible location where this *bona fide* regulation could take place is in the T-zone of lymph nodes, LNs (**Fig 3A**), where T-cells become activated yet must refrain from secreting inflammatory cytokines to avoid damaging the host organ. While LNs are known to be hypoxic which could affect T cell activity(*20*), measurements of LN pH are lacking. Paracortical regions stain positive CD3-T-cells, and manifest some degree of regional hypoxia with pimonidazole staining (**Fig 3B**). LN pHe can be probed using pH indicators injected into either the tail vein or footpad (**Fig S4A**). The pH Low Insertion Peptide, pHLIP^®^ (*21, 22*), conjugated to an 800 nm-emitting fluorescent dye (IR800), was injected into the footpad (40-60 μl of 40 μM), and LNs were harvested 24 hr later for *ex vivo* imaging. Far-red emitting dyes minimize interference from autofluorescence in complex tissues, such as LNs. Paracortical regions were fluorescent, indicating compartments of low pHe (**Fig 3C**). More quantitative measurements of LN pHe were made using 70kDa-dextran conjugated cSNARF1(*23*), i.e. a derivative that cannot passively cross cell membranes, and therefore reports pHe. cSNARF1 excitation (514 nm) and emission (590-640nm) are outside the range of autofluorescence, and the background signal was minimal (**Fig S4B**), thus permitting ratiometric analyses of pH. Dye was injected into the tail vein or the footpad, and measured by intravital imaging (IVI) in anesthetized mice. Fluorescence was collected with a 10X dry objective (NA=0.4), and the ratio was converted offline to pH using a calibration curve determined in saline (**Fig S4C**). Optimal IVI images were obtained 1 hr after tail vein injection or 2-3 hours after footpad injection. To cover an adequate area of the LN, images were acquired in an overlapping cascade of fields-of-view (**Fig S4D**) using a pipeline summarised in **Fig S4E**. Conveniently, tail vein injection allowed for simultaneous measurements of pHe in blood vessels that could serve as a reference for alkaline pHe. LN pHe maps all showed distinct areas of profound acidity in paracortical areas (**Fig 3D**). Experiments were also performed on mice that had been injected with lipopolysaccharide (LPS) to enlarge LNs, or had been treated with oral bicarbonate to test whether raised systemic buffering could neutralize LN acidity (**Fig 3D**). The intensity histograms of pHe values within the LN boundary were analysed by mixed Gaussian modelling to determine the number of compartments (**Fig 3E**): three compartments were identified with tail vein-injected cSNARF1, whereas footpad injections produced a two-compartment distribution. In both preparations, the most acidic LN compartment had a mean pHe of ~6.3, and was surrounded by a compartment of pHe ~6.7 (**Fig 3F**). The most alkaline component, corresponding to blood vessels detected with tail vein delivery, had a mean pHe of 7.1, i.e. as expected from venous blood emerging from an acidic organ. There was no significant effect on LN pHe of treatment with LPS or bicarbonate (**Fig 3F**). Enhanced CO_2_/HCO_3_^-^ buffering would be expected to restore alkaline pH in compartments that are acidic because of diffusive restrictions(*24*), and the finding that oral bicarbonate did not affect LN pHe argues that the source of acidity is likely an active epithelial transport process, equipped with a good feedback mechanism to maintain pHe at desirably acidic set-points. Indeed, and emerging view of lymph node epithelia is that it is a dynamic, active participant in adaptive immunity(*25, 26*).

**Figure 3:**
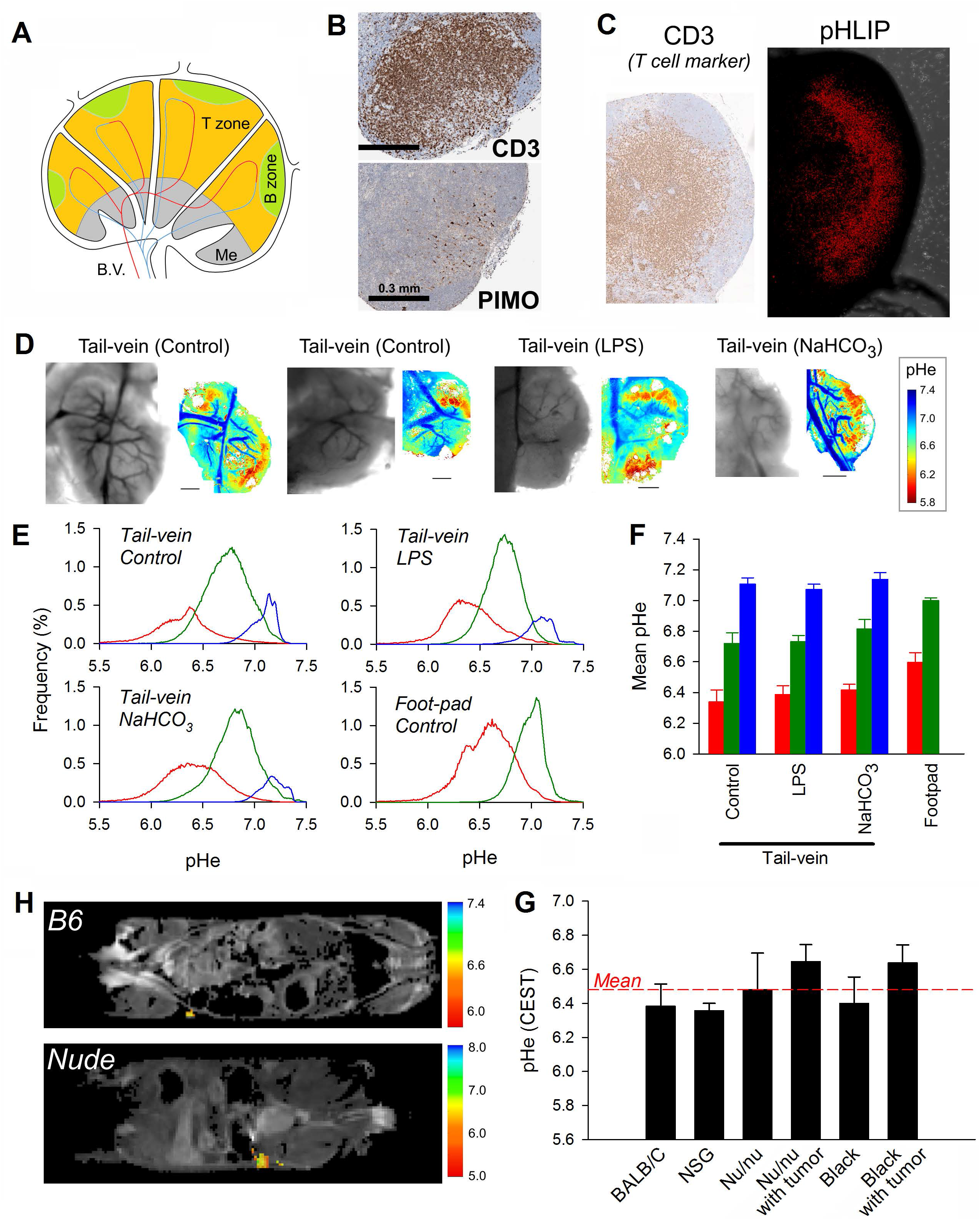
Lymph nodes are acidic. (A) Cartoon of lymph node (LN) showing areas occupied by T- and B-cells, blood vessels (B.V.) and the medulla (Me). (B) Inguinal LN staining for hypoxia. Pimonidazole (Pimo), injected to mice i.p. (60 mg/kg in 100μl) prior to euthanizing, shows punctate staining in the T-zone only, identified by staining for T-cell marker CD3. (C) Fluorescence of pHLIP IR800 (40 μM in 60 μl), injected to mice f.p. prior to euthanizing, was detected in the T-zone only, suggesting an acidic compartment. (D) Intravital imaging of pH-sensitive cSNARF1 fluorescence in inguinal LN. Mice were injected with dextran-conjugated cSNARF1 into the tail vein (20 mg/ml in 100 μl). Measurements on control mice, or mice treated with LPS or on bicarbonate (NaHCO_3_) therapy. (E) Statistical distribution of pHe data analysed by Gaussian mixed models to separate pixels into clusters, representing compartments. Each plot shows the pH-distribution in each of the LN compartments, averaged for all LNs measured for a given experimental class. Note that compared to foot-pad injections, tail vein injections detect an additional compartment corresponding to blood vessels. (F) Summary data for each LN compartment. Mean±SEM of 4, 3, 4, 3 LNs. (H) CEST pH Maps on control (B6) and nude (*nu/nu*) mice, injected with Isovue^®^. pHe maps over inguinal LN region-of-interest (ROI) are overlaid on anatomical T2-weighted images and non-ROI data were zeroed. Heat-bars show the pH scale in each case. (G) The bar graph of Mean ± SEM pHe measured in different animal cohorts (6, 3, 4, 4, 4, 5 mice).

Because IVI may also have artefacts from the surgery used to install the optical window, we also measured LN pHe non-invasively using chemical exchange saturation transfer (CEST) MRI(*27, 28*). In this technique, MR images are collected from different saturation frequencies in mice injected with the CT contrast agent, iopamidol (Isovue^®^, **Fig. S5A**), which has two ionizable groups that resonate at different frequencies that can be interrogated with frequency specific excitations (**Fig. S5B, C**). These resonances have different pH profiles (**Fig S6A-C**). Thus, ratios of saturation occurring at two different frequencies can be used to report pHe (**Fig S6D**). pHe in the LN was mapped in BL/6 mice (**Fig 3G,H**), showing an average pH of 6.38±0.36 (n=9). This acidity was unlikely to be entirely a consequence of residing T-cells because a comparable pHe (6.48±0.43) was reported in *nu/nu* mice (**Fig 3G, S6C**). Also, low LN pHe did not appear to depend on cytokine elaboration, as the same value (6.34±0.34) was measured in NSG mice (**Fig. 3G, S6B**). Further, the LN pHe was no different in B6 mice bearing Panc02 tumours (6.68±0.26), or NSG mice bearing PC3 (6.64±0.20) tumours. These results are concordant with fluorescence-based measurements.

Prior reports have indicated that acid pH can actually oromote dendritic cell (DC) antigen presenting activity(*29, 30*). To evaluate if LN acidity affects T-cell activation, the *in vivo* process was modelled *in vitro* by co-culturing T-cells with dendritic cells (DCs) and antigen (OVA) at pHe 6.6 or 7.4. After a 24-hour activation period, cells were transferred to fresh media at pHe 7.4 to simulate T-cell migration out of the acidic LN towards alkaline tissues, where effector functions are to be exercised (**Fig. 4A**). Monocyte-derived DCs efficiently took-up FITC-tagged OVA protein (**Fig 4B**) and presented OVAsiinfekl peptide (**Fig 4C**) when cultured *in vitro* at either pHe 6.6 or 7.4, indicating that acidic pH did not impair the ability of DCs to process and present antigen to T-cells. No differences in the expression of the DC marker CD40 were seen in cells cultured at pHe 6.6 or 7.4 (**Fig 4D**). Thus, any pHe-dependent effects would be attributable to responses intrinsic to T-cells, rather than a result of insufficient stimulation by DCs. Relative to the control experiment at 7.4, T-cells activated by DCs at pHe 6.6 had decreased IFNγ production measured 24 hours after primary activation (**Fig. 4E**). In contrast, T-cells that had been activated at pHe 6.6, and then rested in pHe 7.4 for 24 hours, produced a comparable level of IFNγ to control cells (**Fig. 4E**). Indeed, IFNγ production from acid-conditioned cells was already restored within 3 hours of incubation at pHe 7.4 (**Fig. 4F**). The restoration of effector function was also observed in T-cells activated with OVAsiinfekl peptide alone (**Fig. S7A**) or with OVAsiinfekl peptide in presence of DCs (**Fig. S7B**). Indeed, the recovery of IFN production moving from pH 6.6 to 7.4 was more robust in cells incubated with DCs and peptide. Thus, T-cell functions were observable within as little as 3 hours of exposure to alkaline conditions, if their prior activation had occurred in the presence of DCs at acidic pHe. This is similar with earlier findings (**Fig 1F**) that, after 24 hr of activation at pH 7.4, T cells could be held static for up to 48 hr by placing them at pH 6.6, and were able to immediately resume IFNγ elaboration when released into an environment at pH 7.4. This relatively fast time course of effector function restoration was contrasted to a control experiment, in which T-cells were first incubated without stimulation at pHe 6.6 or 7.4 for 24 hours, and then subsequently activated with OVAsiinfekl peptide under alkaline conditions (**Fig. S8A**). Cells from both incubation conditions produced similar levels of IFNγ at both 3 and 24 hours following simulation, measured by ELISA of media (**Fig. S8B**), or by flow (**Fig. S8C**). Collectively, these findings indicate that T-cells presented with antigen by DCs at low pHe become activated yet remain inactive, and are able to regain effector functions within hours of exposure to alkaline pHe. The period in low pHe appeared to hasten the rate at which T-cells restore their effector functions in alkaline conditions.

**Figure 4:**
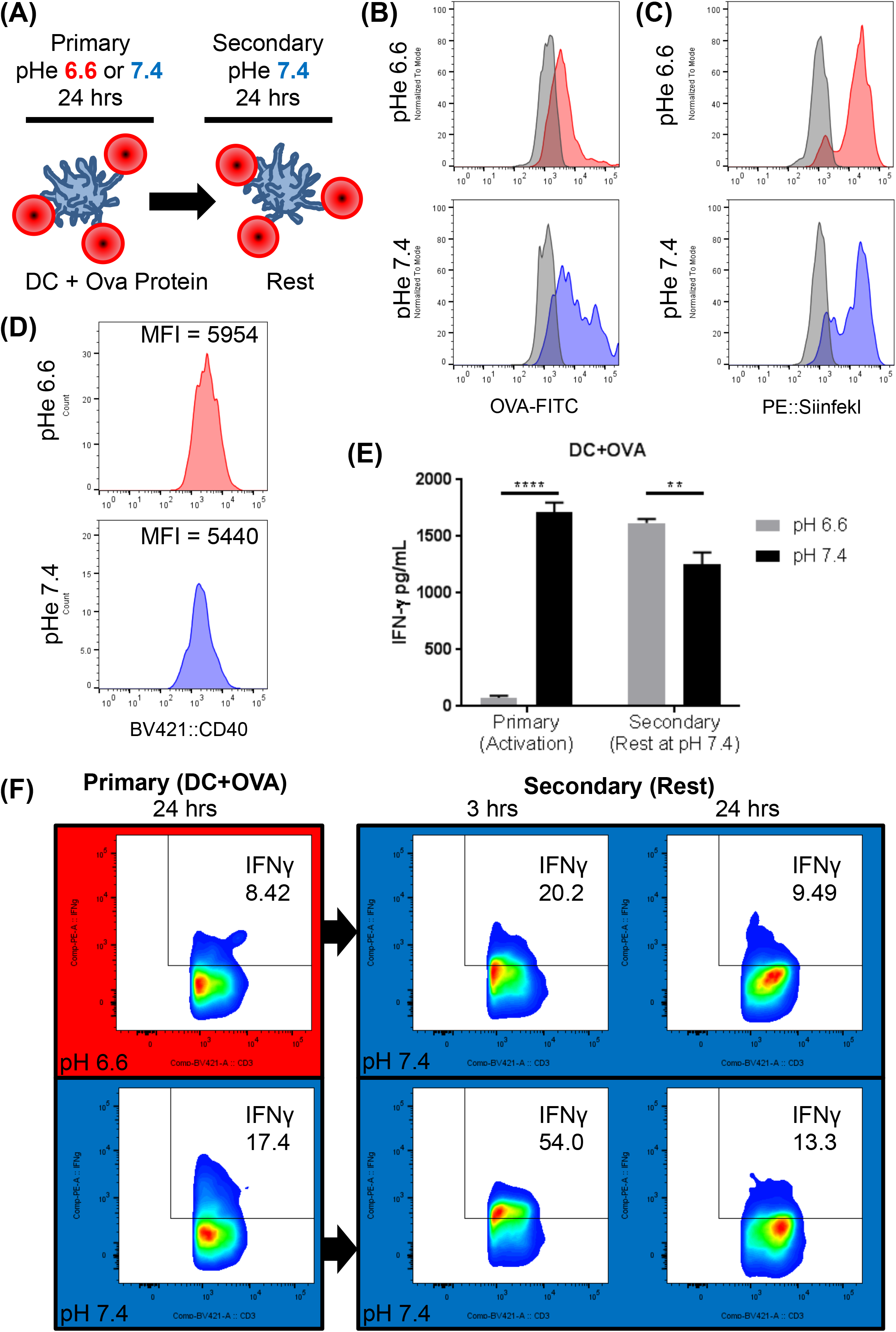
Rescue of low pHe effects on IFN-γ production. (**A**) The effects of pHe on T-cell activation was modeled by activating T-cells with DC presenting OVA protein peptides at pHe 6.6 or 7.4. Following 24 hours, cells were then transferred to media at pHe 7.4. (**B**) Dendritic cells incubated at pHe 6.6 or 7.4 for 24 hours take up OVA-FITC protein, (**C**), present SIINFEKL peptide, and (**D**) maintain CD40 expression. (**E**) ELISA of IFN-γ production from T-cells activated at pHe 6.6 or 7.4 for 24 hours and then transferred to pHe 7.4 for an additional 24 hrs. (**F**) Intracellular IFN-γ staining of T-cells activated at pHe 6.6 or 7.4 for 24 hours and then transferred to media at pHe 7.4 for an additional 24 hours.

In considering the metabolic reprogamming that occurs during immune evasion, it has been noted by others that “the role of extracellular acidosis is not clearly immune-suppressive, but can have both promoting and suppressive effects on different classes of immune cells”(*31*). Indeed, we documented herein a physiologically beneficial role of acidosis in adaptive immunity wherein paracortical LN acidosis inhibits MCT activity and reduces the glycolytic rate of resident T-cells, supressing their effector fuctions. This mechanism serves as a biological checkpoint that prevents damaging, pro-inflammatory effector functions from being elicited within the LN. Once T-cells migrate out of the LN to alkaline tissues, their effector functions are rapidly activated. Thus, LN acidity may be dually beneficial in normal physiology: by priming T-cells for a faster activation outside the organ, and keeping their effector functions suppressed whilst in residence. It appears however that this mechanism has been co-opted in cancers to evade immune surveillance.

## Supporting information

Supplemental Methods and Data for Wu et al

