## Supplemental Methods and Data for Wu et al for "Lymph Nodes Inhibit T-cell Effector Functions Locally by Establishing Acidic Niches"

#### Materials and Methods

***Isolation and activation of T cells.***

Female B6 (C57BL/6), Pmel, OT-I,OT-II and TDAG8 knockout (TDAG8 KO) mice on the C57BL/6 background were bred and housed at the Animal Research Facility of the H. Lee Moffitt Cancer Center and Research Institute (Tampa, FL). Naïve T cells were isolate from mice spleen using T cell column (R&D system). T cells from B6 mice and TDAG8 knockout mice were cultured in complete medium with 5mg/mL plate-bounded anti-CD3 antibody and 5mg/mL soluble anti-CD28 antibody for 48 hours. Isolated Pmel, OT-I and OT-II T cells are cultured in complete medium with 5μg/mL gp100 $_{25\text{-}33}$ peptide, 10μg/mL OVA $_{\text{SIINFEKL}}$ peptide and 10μg/mL OVA $_{323\text{-}339}$ peptide, respectively. All animal experiments were approved by the Institutional Animal Care and Use Committee and performed in accordance with the U.S. Public Health Service Policy and National Research Council Guidelines.

***Cell lines.*** Jurkat cell was maintained in RPMI-1640 medium with 5% FBS. Jurkat cells were stimulated with PMA and calcium ionophore A31111 for 24 hours. The supernatants were collected for IL-2 quantification by ELISA (BD Biosciences). PC3M, human adenocarcinoma prostate cancer (Caliper) cells, were maintained in MEM/EBSS medium with 5% FBS. $10^6$ cells were subcutaneously injected into right flank of C57BL/6 mice. The isogenic Panc02 pancreatic cancer cells (a kind gift from Emmanuel Zervos, East Carolina University, Greenville, NC) were maintained in RPMI 1640 with 10% FBS. $10^6$ cells were subcutaneously injected into right flank of *nu/nu* mice. Tramp-C2 cells are male derived prostate cancer cells that were maintained in DMEM high glucose medium with 10% FBS. 4 x $10^6$ cells were subcutaneously injected into right flank of C57BL/6 mice.

***Animals**:* All animals were maintained under Institutional Animal Care and Use Committee (IACUC) at H. Lee Moffitt Cancer Center. Eight-to ten-week old Balb/c,

C57BL/6, NSG, and *nu/nu* mice (male, 22-25 g) were purchased from The Jackson Laboratory and used as control mice and as host for TRAMP-C2, Panc02 and PC3 cells.

***Seahorse measurements of metabolism.*** Extracellular acidification rate (ECAR) and oxygen consumption rate (OCR) were measured by Seahorse XF96 Analyzer (Agilent). Cells were cultured with bicarbonate-free RPMI-1640 medium with 2 mM HEPES and 2 mM MES. The buffering capacity was determined to calculate the proton production rate (PPR).

***Flow cytometry.*** Fresh isolated T cells were activated at pHe 7.4 or pHe 6.7 for 72 hours. Cells were collected and wash by PBS twice, then stained in FACS buffer with the following antibodies for flow cytometric analysis: CD3, CD4, CD8, CD44 and CD62L (all from BD Biosciences). Live/Dead fixable near-IR (Invitrogen) was used to extrude dead cells before analysis. To analyse intracellular marker IFNγ, cells were stained with surface marker and Live/Dead dye, fixed and permeabilized by Fixation/Permeabilization Solution Kit (BD Biosciences), and then stained with anti-IFNγ antibody. Samples are analysed by LSR II Flow Cytometer (BD Biosciences).

***Antibodies.*** Anti-pimonidazole antibody (#PAb2627, a rabbit polyclonal antibody) was purchased from Hyproxyprobe, Inc (Burlington, MA) and used at a 1:100 dilution; anti-CD3 antibody (#M3072, a rabbit monoclonal antibody) was purchased from Spring Bioscience Corp. (Pleasanton, CA) and used at a 1:100 dilution.

***Cytokine Beads Array assay.*** T cells were activated for 48 hours and re-stimulated at pHe 7.4 or pHe 6.7 for 24 hours. Culture medium was collected for cytokine beads array analysis according to the manufacturer's manual (BioLegend). Briefly, 25μL culture medium was sequentially mixed with antibody-conjugated beads, detection antibody and SA-PE. Washed samples were analysed by flow cytometer.

***Cell proliferation assay.*** Fresh prepared T cells were washed by PBS twice and stained with 2 uM CellTrace Voilet in PBS for 10 minutes, and incubated in complete medium for another 20 minutes to quench residual dye. After two wash with complete medium, cells were activated at pHe 7.4 or pHe 6.7 for 72 hours. After activation, cells were collected and stained with surface marker and live/dead dye for before analysis.

***Immunochemistry.*** Murine inguinal lymph nodes were surgically removed from C57Bl/6 mice, fixed in formalin and paraffin embedded. Slides were prepared with 4 μm thick tissue slices and stained using a Ventana Discovery XT automated system (Ventana Medical Systems) as per manufacturer's protocol with proprietary reagents. The primary antibodies were used to detect pimonidazole (1:100, Hyproxyprobe #PAb2627) and CD3 (1:100, Spring Bioscience #M3072) expression. Slides were incubated with Ventana OmniMap Secondary Antibody followed by Ventana ChromoMap kit to detect the proteins staining and then slides were counterstained with Hematoxylin.

***Superfusion.*** Superfusion experiments were performed in a plastic chamber (Pecon, TempController 2000-2) supplied by a solution line with a switcher that changed between one of two lines (the other being diverted to the waste bottle). The plastic chamber was mounted on a confocal microscope and heated to 37°C by small scale temperature incubator (The Cube Life Imaging Services). Solution exchange was attained with a time constant of 2.6 s. Solution flows were 2-4 ml/min.

***Solutions and media.*** (i) Solutions for seahorse experiments: 2 mM HEPES, 2 mM MES, 5.3 mM KCl, 5.6 mM Na-Phosphate, 11 mM glucose, 133 mM NaCl, 0.4 mM $MgCl_2$, 0.42 mM $CaCl_2$, titrated to given pH with NaOH. For reduced $Cl^-$ experiments, 133 mM NaCl was replaced with 133 Na-Gluconate and $MgCl_2$ and $CaCl_2$ were raised to 0.74 and 1.46 mM, respectively, to account for gluconate-divalent binding. Amount of dilute HCl or NaOH added to medium to reduce pH to target level was determined empirically. Solutions for pH measurements under superfusion: For pH 7.4, 133 mM NaCl, 5.3 mM KCl, 10 mM Glucose, 1 mM $CaCl_2$, 1 mM $MgCl_2$, 22 mM $NaHCO_3$. For lower pH, $NaHCO_3$ was reduced (compensated by NaCl) to attain a target pH, according to the Henderson Hasselbalch equation ($pH = 6.15 + \log([HCO_3^-]/[CO_2])$, where $[CO_2]$ is 1.2 mM for 5%. All solutions were bubbled in 5% $CO_2$. (iii) Solutions for $Ca^{2+}$ imaging: For pH 7.4, 133 mM NaCl, 5.3 mM KCl, 0.8 mM $MgCl_2$, 0.9 mM Na-Phosphate, 22 mM $NaHCO_3$ and either 1.8 mM $CaCl_2$, 0.5 mM $CaCl_2$ or 0.5 mM EGTA. For pH 6.6, $NaHCO_3$ was reduced to 2.75 mM and NaCl raised accordingly. All solutions were bubbled with 5% $CO_2$/balanced air. (iv) Calibration solutions for nigericin: 145 mM KCl, 1 mM $MgCl_2$, 0.5 mM EGTA, 10 mM HEPES, 10 mM MES and pH adjusted with NaOH to required level.

***Confocal imaging.*** Imaging was performed on an SP5 system (Leica Microsystems). Cellular measurements were performed with an oil-immersion x63 objective and intravital microscopy was performed with a dry x1.6 or x10 objective. The following excitation (ex) and emission (em) wavelengths were used: dextran-conjugated cSNARF1 (cat#D3304, Invitrogen): 514 nm ex, 580/640 nm em (during lymph node imaging); Hoechst 34580 (cat#H21486, Thermo-Fisher): 405 nm ex, 420 nm; FuraRed: 488 nm ex, 585/685 nm; pHLIP IR800 (was a gift from the Dr. Yana Reshetnyak's lab, University of Rhode Island): 633nm ex, >700 nm. Settings were optimised to obtain maximal quality under the constraints of temporal resolution.

***Lymph Node Imaging:*** The inguinal lymph node is exposed for imaging on the confocal microscope by performing a midline incision to separate the skin from the peritoneum. The skin is pinned down and the excess fat around the inguinal lymph node is carefully removed with sterile Dumont #5 forceps. Once cleared and exposed, a 3D printed window chamber, with a 12 mm in diameter window, is placed over the lymph node area. The chamber is secured using a tissue adhesive (3M Vetbond #1469SB) and 12 mm Micro coverslip (cat# 72226-01 Electron microscopy sciences). We ensure that the lymph node region underneath the coverslip does not dryout by injecting 200 µl of 1X PBS. Mice were kept under anaesthesia (1.5% isofluorane) throughout the surgical procedure and in a 37°C warming chamber during imaging. To measure pH in the inguinal lymph node, mice were injected with dextran-conjugated cSNARF1 (cat# D3304, Invitrogen) at a concentration of 20 mg/ml in 100 µl via their tail vein or foot pad. To determine whether inflammation or buffering would change the pH of the lymph node microenvironment, mice were treated with lipopolysaccharide (LPS, cat# L3012, sigma Aldrich) at a concentration of 1000 ng/kg (i.p. injection) for 48 h or provided mice with 400 mM $NaHCO_3$ *ad libitum* for 9-10 days, respectively, before imaging the mice with dextran-conjugated cSNARF1.

***Image analysis***. Cytoplasmic pH was measured by gating pixels according to a threshold level of Hoechst signal within cSNARF1-positive pixels. Fluorescence at 580 and 640 nm was averaged, background offset and ratioed for each particle representing a cell. For time course experiments, pHi was probed in the entire cell to allow for faster acquisition rates. Buffering capacity was measured from the change in weak acid/base concentration, assuming passive equilibration, and the pH change. Transmembrane acid-base fluxes were therefore calculated as the product of pH change and buffering capacity. Cell surface area/volume ratio assumed spherical symmetry i.e. 3/radius. For intravital microscopy, image montages were constructed with in-house software that aligned fluorescence or anatomical landmarks.

***CEST imaging.*** Magnetic resonance data were acquired with a 7 T horizontal magnet Bruker equipped with nested 205/120/HDS gradient insert and a bore size of 310 mm. A 35 mm Litzcage coil (Doty Scientific) was used to carry out all experiments. Before imaging, animals were placed in an induction chamber and anesthetized with 3% isoflurane delivered in 1.5 liter/min oxygen ventilation. After complete induction, animals were restrained in a custom-designed holder and inserted into the magnet while constantly receiving isoflurane (1 to 3%) within the 0.6 liter/min oxygen ventilation. Body temperature (37° ± 1°C) and respiratory functions were monitored continuously (SAII 177 System) during the experimental time. Coronal T2-weighted fast spin-echo multislice experiments (acquired with TE/TR [echo time/repetition time] = 31 ms/2271 ms, field of view (FOV) = 80 × 30 $mm^2$, matrix = 256 × 96, yielding a spatial in-plane resolution of 312 µm, slice thickness of 1.5 mm). That image was used as anatomical images for the pH map. CEST images was done using an optimized version of one published previously(*1*); TE/TR = 10 ms/10 s, the saturation used was 3 µT during 5 s, same FOV than T2 but matrix = 171 x 64. Only one slide was used in CEST, the one containing the inguinal lymph node. Animals were injected with ISOVUE370 (Bracco) at

300 ul iv bolus injection; followed by an iv infusion of 300 μl/h. To create the CEST maps, an in-lab designed Matlab code was used. The pH values were calculated after a calibration curve done in the same system with 20 mM ISOVUE370 phantoms.

#### Legends for Supplemental Figures

**Figure S1:** (A) Relationship between cytokine levels measured in paired experiments at low and high pH by Cytokine Beads Array assay, showing reduced release in acidic conditions for most cytokines, with the exception of those shown in purple. Data is showed as Mean±SD. (B) Interferon-γ (IFNγ) immunoreactivity is stable over a range of pHe. Data is showed as Mean±SD. (C) Low pHe polarizes mitochondrial membrane potential of T-cell measured by confocal imaging and (D) confirmed by flow cytometry. (E) Proliferation rate of $CD4^+$ and $CD8^+$ cell from total B6 T cell at low and high pH. (F) Fraction of $CD44^+CD62L^+$ cells (memory phenotype) is increased at low pHe in CD4+ and CD8+ subpopulation from B6 T cell.

**Figure S2:** Investigating methods of rescuing acid-inhibited IFNγ release. (A) G-protein-coupled receptor protein TDAG8 homogeneous ($TDAG^{-/-}$) and heterogeneous ($TDAG8^{-/+}$) knockout and OGR1 knockout ($OGR^{-/-}$) showed not rescue of IFNγ production at low pHe. (B) OGR1 inhibitor BA-39-PQ30 and (C) GPR4 inhibitor NE-52-QQ57 did not rescue IFNγ production at low pHe. (D) Inhibitors of TRPV1, (E/F) ASIC-3, (G) ASIC1a, (H) ENaC and (I) NHE1 were unable to rescue IFNγ production at low pHe. (J) The OGR1-selective positive allosteric modulator OGERIN did not rescue the IFNγ release at low pHe. (K) The phorbol 12-myristate 13-acetate plus A23187 and (L) a low dose of Trichostatin A, a histone deacetylase inhibitor, partially restored IFNγ production at pHe 6.6, compared to pHe 7.4. (M) A low dose of Trichostatin A increased the ratio of IFNγ production at pHe 6.7 and pHe 7.4. (N) The measurement of pHi by flow cytometry, left, calibration curve; right, pHi signal of cells under high and low pHe. Data is expressed as Mean±SD, experiment is repeated at least twice.*, $p<0.05$.

**Figure S3:** *Investigating the mechanism of acid-inhibition of T-cell functions*. (A) Protocol for interrogating store-operated $Ca^{2+}$ entry. Superfused OT-I T-cells were loaded with FuraRed and exposed to 10 μM thapsigargin for 10 min in $Ca^{2+}$-free (0.5 mM EGTA) conditions to deplete the endoplasmic reticulum (ER) $Ca^{2+}$ store. Next, extracellular $Ca^{2+}$ was raised to 0.5 mM by rapid solution switching at either pHe 7.4 or 6.6. The slope and extent of intracellular $Ca^{2+}$ rise (FuraRed ratio) is a readout of $Ca^{2+}$ entry. Ionomycin (10 μM) was added at the end of the experiment as a positive control for $Ca^{2+}$ entry. (B) Experiment repeated with higher $Ca^{2+}$ (1.8 mM) in OT-I cells. (C) Experiment repeated with B6 cells using 1.8 mM $Ca^{2+}$. Time courses show mean±SEM of 15-25 cells each. (D) Calibration curve for cSNARF1 AM-loaded into B6 cells, performed by the nigericin (10 μM) method, showing best fit to Grinkiewicz equation. Mean±SEM of results from 4 experiments, each with 40-60 cells each. (E) Phosphofructokinase (PFK) activity is sensitive to pH change. Cell lysate from activated B6 cells are subjected for enzyme activity assay. Mean±SD; **, $p<0.01$; ***, $p<0.001$, compared to pH 7.4. (F) Effect of MCT1 inhibitors AR-C (AR-C155858; 10 μM) and SR (SR13800; 10 μM) on rate of pHi change upon rapid removal of extracellular lactate. The ablated rate arises from inhibited MCT activity. Buffering capacity was assumed to be constant (control). (G) EACR measured by Seahorse, showing effect of adding L- and D-lactate on glycolytic rate. Note that due to the stereo-specificity of LDH, the inhibitory effect of the L-isoform is stronger.

**Figure S4:** *Pipeline for measuring lymph node pH by intravital imaging.* (A) Bright field image of LN in control mice injected with Hoechst-33342 into the tail vein (10 mg/kg in 100 μl) to visualize blood vessels in LN and TexasRed dextran (50mg/ml in 100 μl) into the footpad of the same animal to demonstrate delivery of substances to the LN. Images collected with x1.6 low power objective. Imaging of LN for Hoechst-33342 (400-440 nm) and TexasRed (600-640 nm) shows uptake of footpad-delivered fluorophore into the LN, relative to its vasculature. Images collected 3 hours after footpad injection. Tail-vein injection occurred 2 hours after footpad injection. (B) Bright field image of LN x1.6 and fluorescence images collected by x10 objective acquired in mice that had not been injected with cSNARF1 conjugated with dextran. Lack of fluorescence signal indicates that autofluorescence was negligible. Note that the same optical and electronic settings have been used as in control experiments. (C) Calibration curve for cSNARF1-conjugated to 70kDa dextran collected with x10 objective, with best fit to Grinkiewicz equation, showing that the dye is optimal for detecting pH in the range 6-8. (D) Bright field image of LN at x1.6 showing areas were fluorescence was acquired with a higher power (x10) objective. The pHe map of the LN was then constructed by merging the overlapping cascade of images, using anatomical or fluorescence landmarks to align images correctly. (E) Pairs of fluorescence at 580 nm (560-600 nm) and 640 nm (620-660 nm) for the four fields of view acquired with x10 objective.

**Figure S5** – Chemical Exchange Saturation Transfer (CEST). (A) Structure of iopamidol (Isovue®) showing ionizable amide groups in red and blue. The two blue groups are co-resonant. (B) Z-spectrum generated by measuring water resonance intensity following saturating frequency specific pulses at between +5 and -5 ppm relative to the water resonance at 0 ppm. The two amide resonances are indicated with red and blue arrows. (C1-3) Images of phantoms at different pH values. (C1) T2-weighted images of all the phantoms, and their saturation intensities at (C2) 4.2 and (C3) 5.5 ppm, respectively.

**Figure S6.** (A) Z-spectra of each phantom; symbols are raw data and lines are data after $B_0$ correction. (B) Saturation profiles of the 4.2 and 5.5 ppm resonances between pH 5.0 and 8.0. (C) Ratio of 4.2 to 5.5 resonances as a function of pH; and (D) the final calibration polynomial best fit curve calculated using the ratiometric approach and the saturation profiles obtained for these phantoms.

**Figure S7, Rescue of low pHe effects on IFN-γ production in T-cells activated with SIINFEKL peptide. (A)** Intracellular IFN-γ staining of T-cells activated at pHe 6.6 or 7.4 for 24 hours with SIINFEKL peptide alone or (**B**) dendritic cells presenting SIINFEKL peptide. Both sets of cells were then transferred to media at pHe 7.4 and IFNγ production measured by flow at 3 and 24 hours thereafter.

**Figure S8. Resting T-cells in pHe 6.6 or 7.4 has no effect on IFN-γ production at pHe 7.4**. (**A**) T-cells were rested at pHe 6.6 or 7.4 for 24 hours and then activated with SIINFEKL peptide at pHe 7.4 for an additional 24 hours. (**B**) IFN-γ production was measured during initial rest period and activation period by ELISA (**C**) and intracellular IFN-γ staining (**C**).


1. D. L. Longo *et al.*, A general MRI-CEST ratiometric approach for pH imaging: demonstration of in vivo pH mapping with iobitridol. *J Am Chem Soc* **136**, 14333-14336 (2014).

| T-cell | Extra-cellular Ion | pHe | pHi | EACR (relative) | EACR (mM/min) | Lactic acid Perm-eability (µm/s) | [Lactate] (mM) |
|---|---|---|---|---|---|---|---|
| B6 | Cl | 6.6 | 6.916 | 0.503 | 5.028 | 1730 | 3.302 |
| B6 | Cl | 6.8 | 6.995 | 0.728 | 7.275 | 2448 | 4.051 |
| B6 | Cl | 7 | 7.027 | 1.006 | 10.057 | 3572 | 4.131 |
| B6 | Cl | 7.2 | 7.077 | 1.272 | 12.718 | 5319 | 3.937 |
| B6 | Cl | 7.4 | 7.092 | 1.492 | 14.921 | 8004 | 3.183 |
| OT1 | Cl | 6.6 | 6.886 | 0.326 | 3.264 | 1730 | 2.002 |
| OT1 | Cl | 6.8 | 6.947 | 0.56 | 5.604 | 2448 | 2.795 |
| OT1 | Cl | 7 | 6.990 | 0.864 | 8.639 | 3572 | 3.263 |
| OT1 | Cl | 7.2 | 7.062 | 1.204 | 12.043 | 5319 | 3.603 |
| OT1 | Cl | 7.4 | 7.088 | 1.525 | 15.248 | 8004 | 3.218 |
| B6 | Glu | 6.6 | 7.075 | 0.715 | 7.147 | 1730 | 6.778 |
| B6 | Glu | 6.8 | 7.136 | 1.061 | 10.61 | 2448 | 8.188 |
| B6 | Glu | 7 | 7.196 | 1.259 | 12.588 | 3572 | 7.646 |
| B6 | Glu | 7.2 | 7.244 | 1.562 | 15.62 | 5319 | 7.114 |
| B6 | Glu | 7.4 | 7.309 | 1.857 | 18.566 | 8004 | 6.528 |
| OT1 | Glu | 6.6 | 7.078 | 0.738 | 7.378 | 1730 | 7.049 |
| OT1 | Glu | 6.8 | 7.185 | 1.093 | 10.929 | 2448 | 9.436 |
| OT1 | Glu | 7 | 7.225 | 1.166 | 11.663 | 3572 | 7.572 |
| OT1 | Glu | 7.2 | 7.269 | 1.464 | 14.643 | 5319 | 7.057 |
| OT1 | Glu | 7.4 | 7.309 | 1.746 | 17.462 | 8004 | 6.129 |

**Table S1**: Estimate of intracellular lactate, based on metabolic rate and transmembrane pH gradient.

### FigureS1

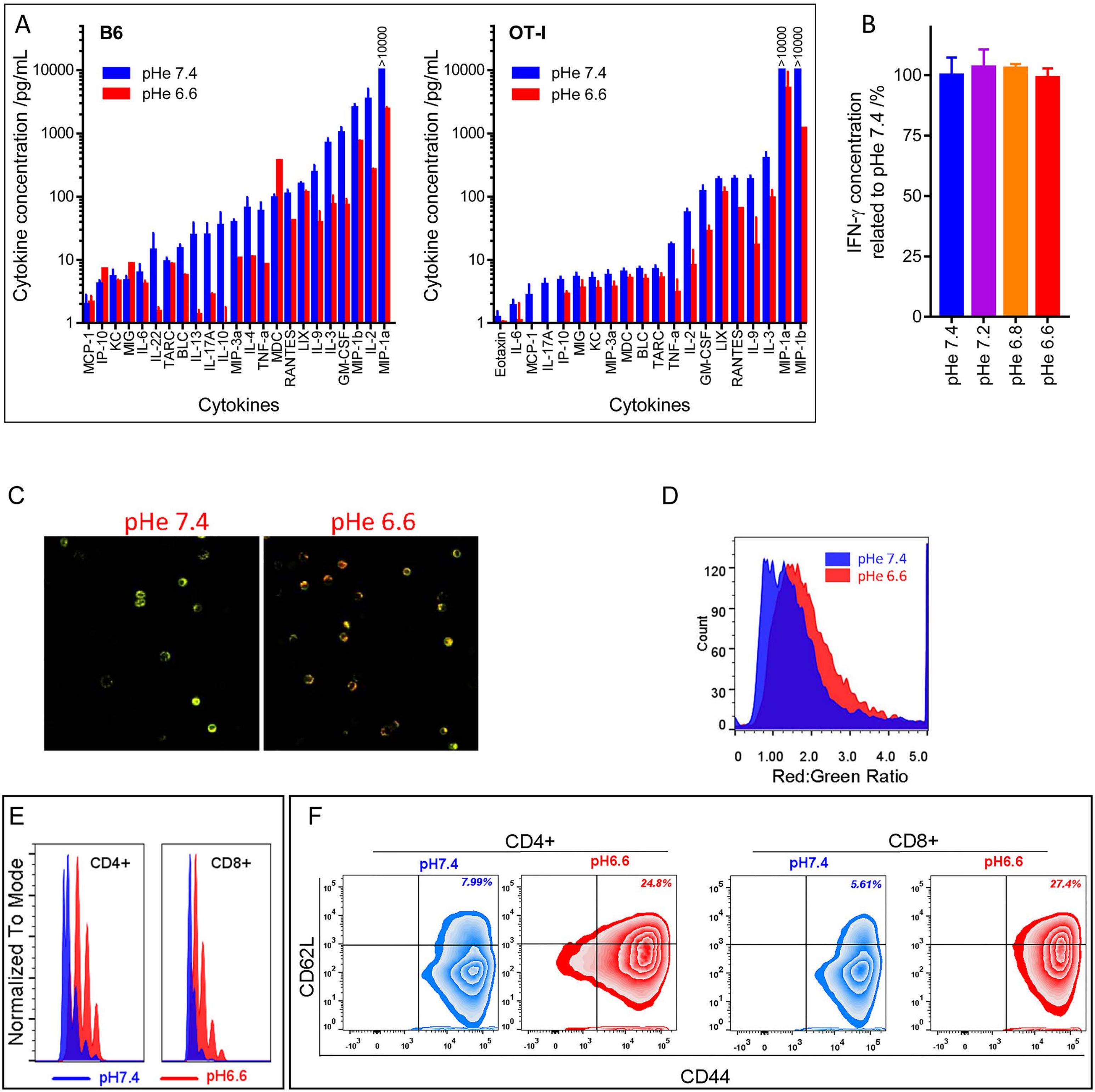

A
B6
pHe 7.4
pHe 6.6
>10000
Cytokine concentration /pg/mL
10000
1000
100
10
1
MCP-1
IP-10
KC
MIG
IL-6
IL-22
TARC
BLC
IL-13
IL-17A
IL-10
MIP-3a
IL-4
TNF-a
MDC
RANTES
LIX
IL-9
IL-3
GM-CSF
MIP-1b
IL-2
MIP-1a
Cytokines
OT-I
pHe 7.4
pHe 6.6
>10000
>10000
Cytokine concentration /pg/mL
Eotaxin
IL-6
MCP-1
IL-17A
IP-10
MIG
KC
MIP-3a
MDC
BLC
TARC
TNF-a
IL-2
GM-CSF
LIX
RANTES
IL-9
IL-3
MIP-1a
MIP-1b
Cytokines
B
IFN-γ concentration related to pHe 7.4 /%
100
75
50
25
0
pHe 7.4
pHe 7.2
pHe 6.8
pHe 6.6
C
pHe 7.4
pHe 6.6
D
pHe 7.4
pHe 6.6
Count
120
90
60
30
0
0
1.00
2.0
3.0
4.0
5.0
Red:Green Ratio
E
Normalized To Mode
CD4+
CD8+
pH7.4
pH6.6
F
CD4+
CD8+
pH7.4
pH6.6
pH7.4
pH6.6
7.99%
24.8%
5.61%
27.4%
CD62L
CD44

### Figure S2

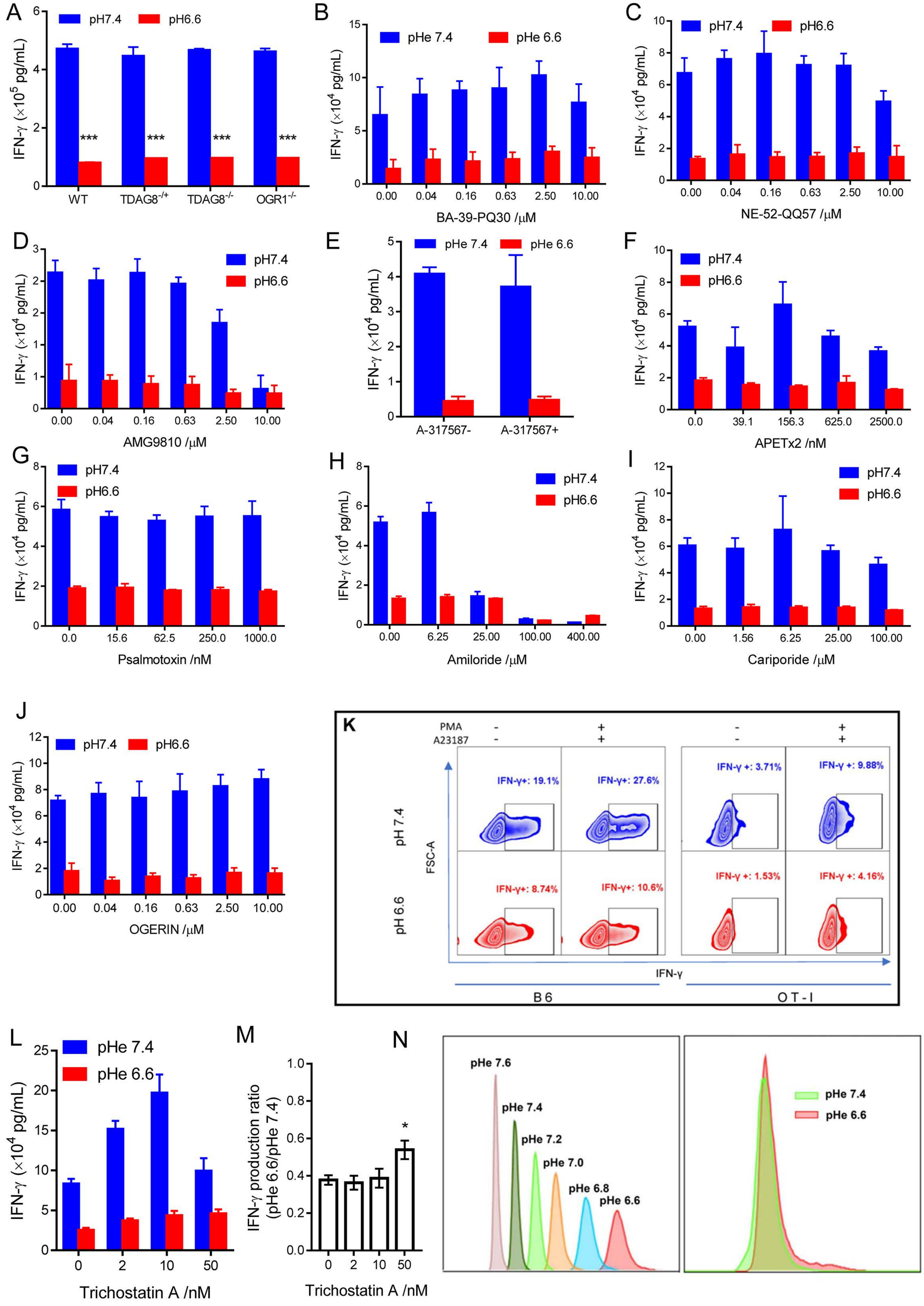
A
pH7.4
pH6.6
IFN-γ (×10^5 pg/mL)
***
WT
TDAG8-/+
TDAG8-/-
OGR1-/-
B
pHe 7.4
pHe 6.6
IFN-γ (×10^4 pg/mL)
BA-39-PQ30 /μM
C
NE-52-QQ57 /μM
D
AMG9810 /μM
E
A-317567-
A-317567+
F
APETx2 /nM
G
Psalmotoxin /nM
H
Amiloride /μM
I
Cariporide /μM
J
OGERIN /μM
K
PMA
A23187
IFN-γ+: 19.1%
IFN-γ+: 27.6%
IFN-γ +: 3.71%
IFN-γ +: 9.88%
IFN-γ+: 8.74%
IFN-γ+: 10.6%
IFN-γ +: 1.53%
IFN-γ +: 4.16%
pH 7.4
pH 6.6
FSC-A
IFN-γ
B6
OT-I
L
Trichostatin A /nM
M
IFN-γ production ratio (pHe 6.6/pHe 7.4)
N
pHe 7.6
pHe 7.4
pHe 7.2
pHe 7.0
pHe 6.8
pHe 6.6

#### FIGURE S3: mechanism

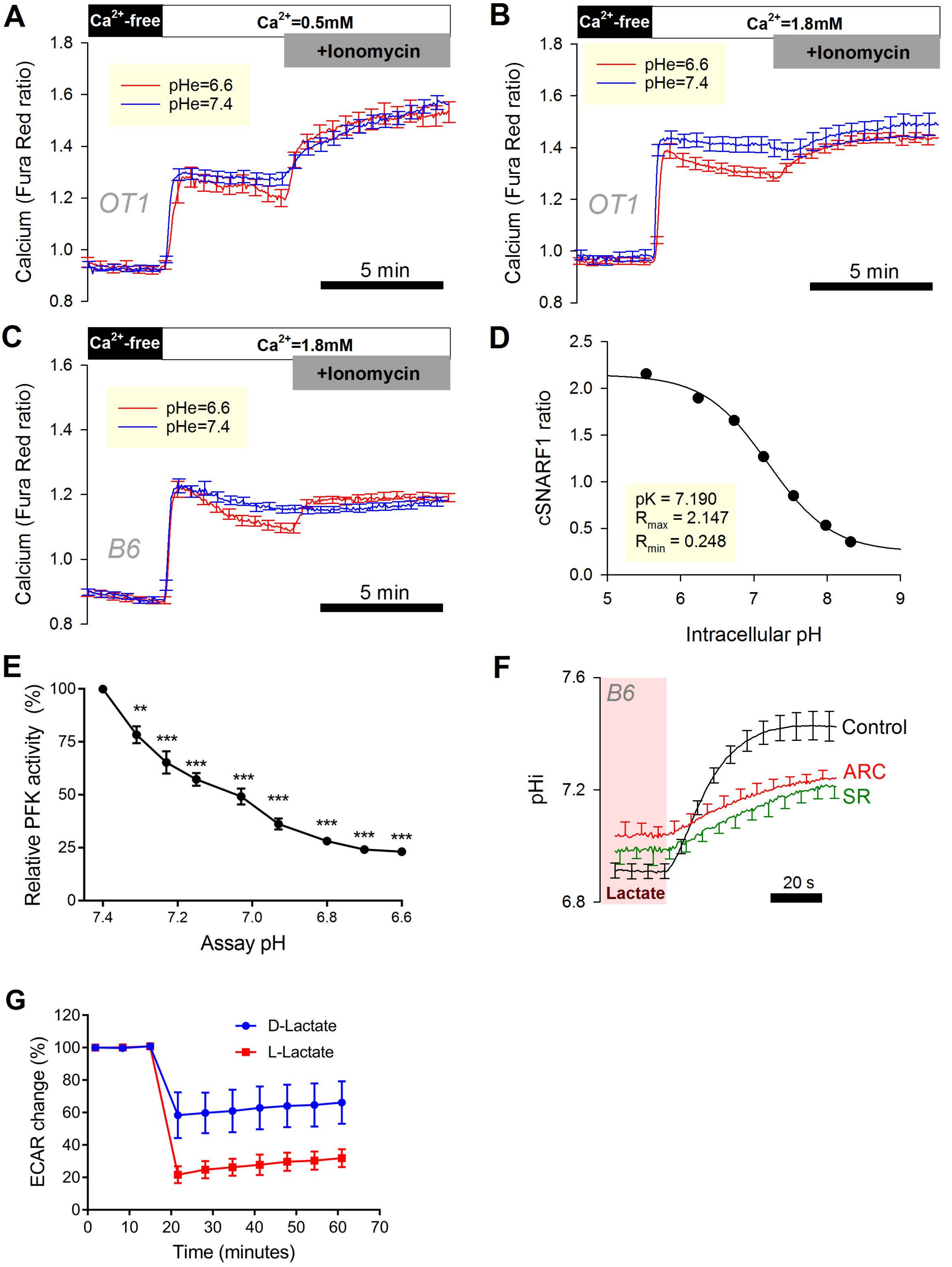

### FIGURE S4: LN pH

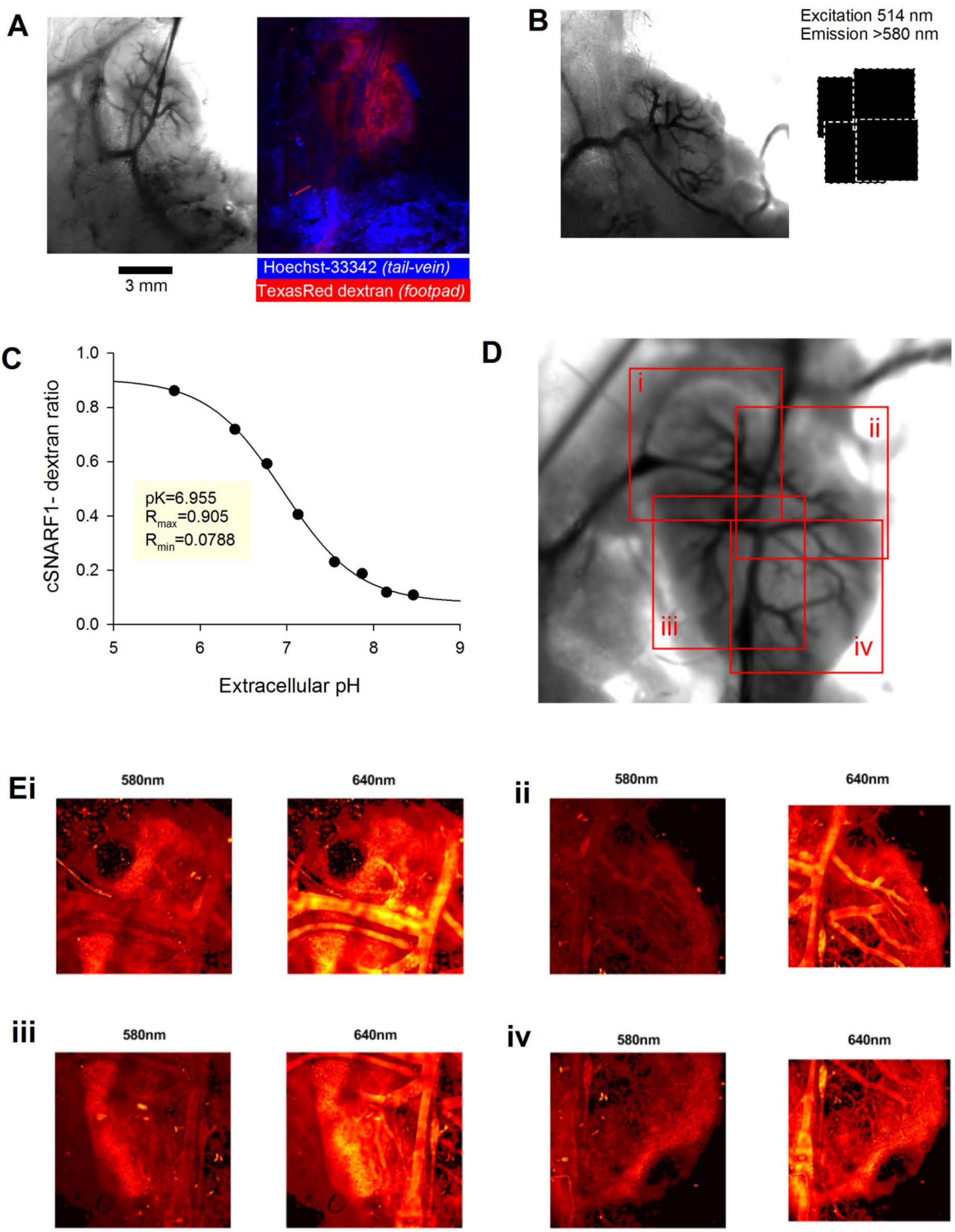

### Figure S5

(A)

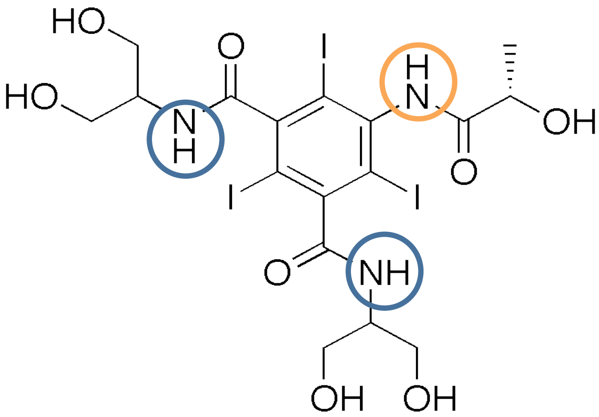


Iopamidol

(B)

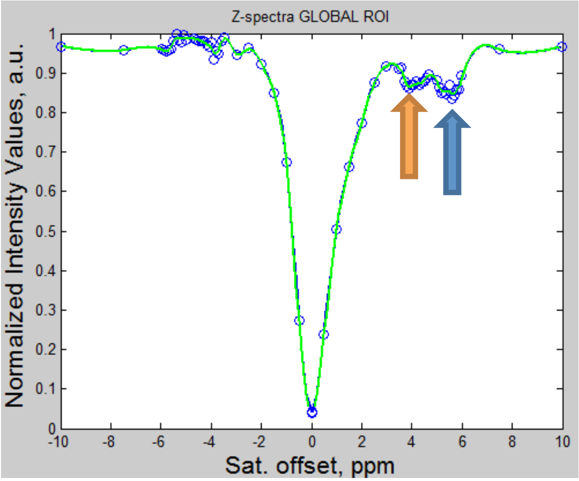


Z-spectrum

(C1)

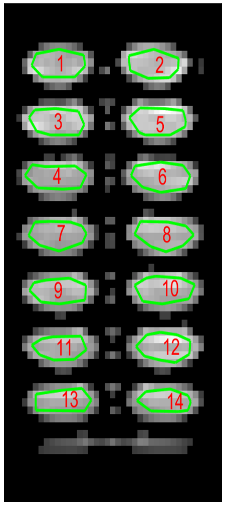


(C2)

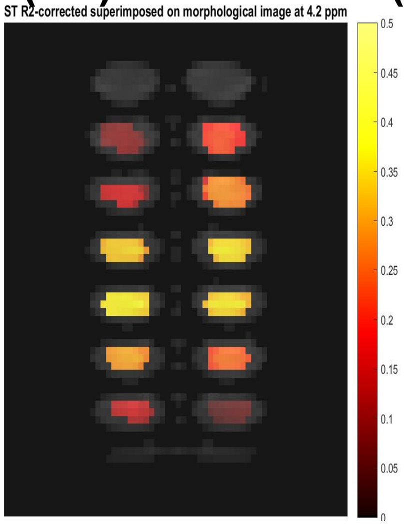


(C3)

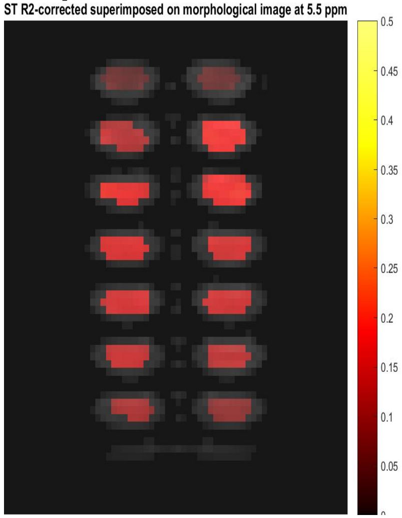


(D)

### Figure S6

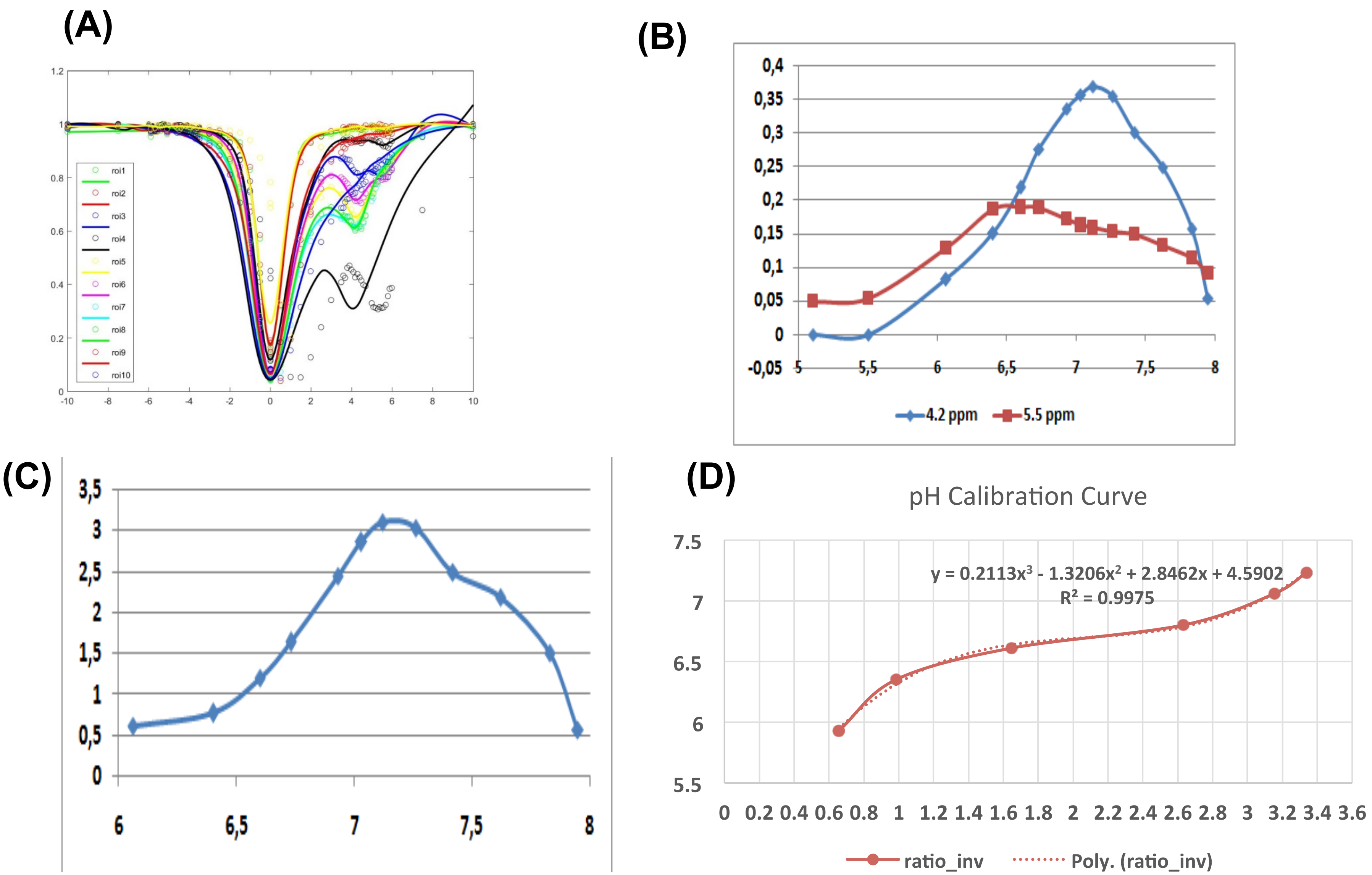

(A)
roi1
roi2
roi3
roi4
roi5
roi6
roi7
roi8
roi9
roi10
(B)
4.2 ppm
5.5 ppm
(C)
(D)
pH Calibration Curve
y = 0.2113x³ - 1.3206x² + 2.8462x + 4.5902
R² = 0.9975
ratio_inv
Poly. (ratio_inv)

**Figure S7**

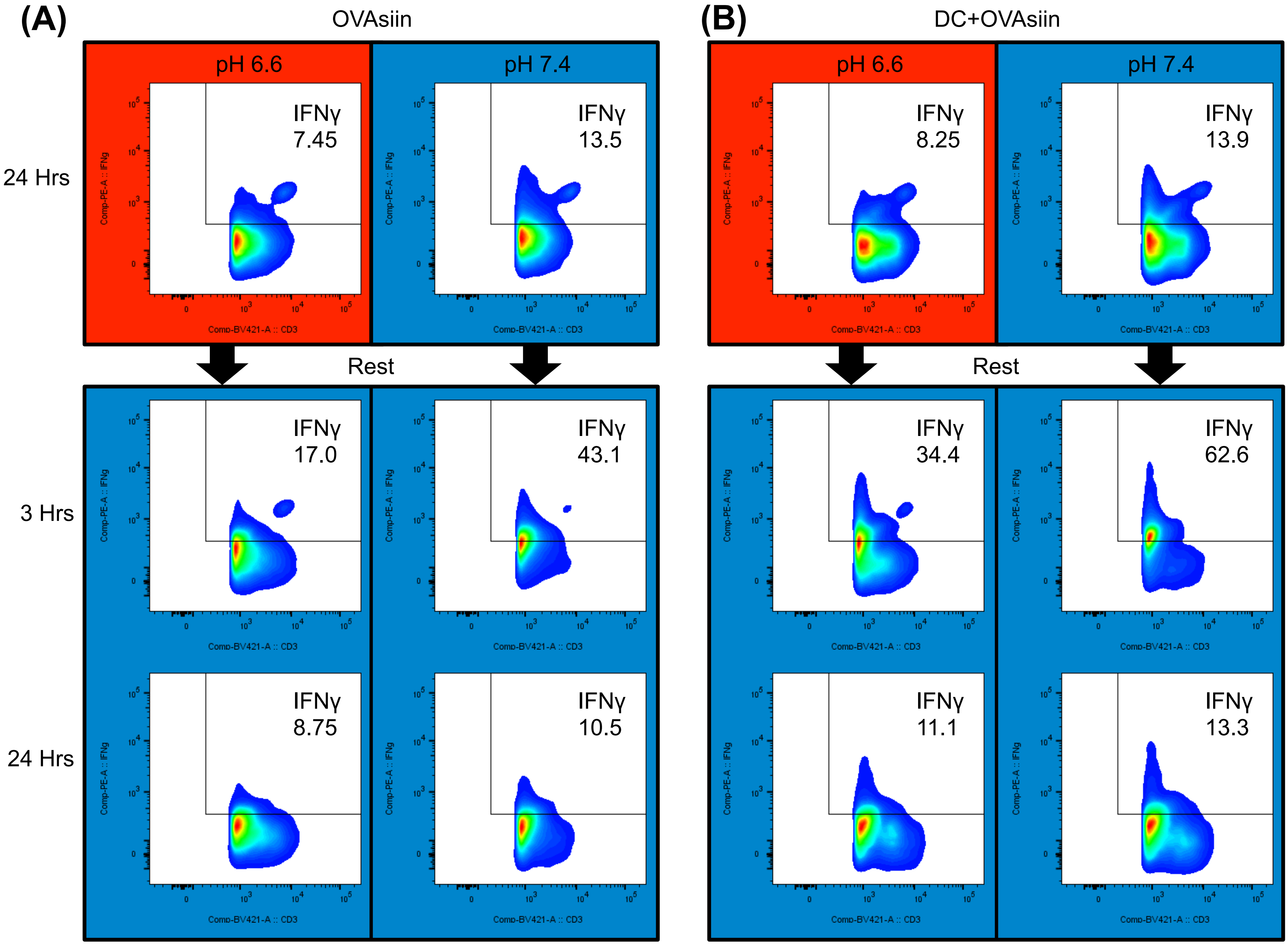
(A)
OVAsiin
(B)
DC+OVAsiin
pH 6.6
pH 7.4
24 Hrs
3 Hrs
24 Hrs
Rest
IFNγ 7.45
IFNγ 13.5
IFNγ 8.25
IFNγ 13.9
IFNγ 17.0
IFNγ 43.1
IFNγ 34.4
IFNγ 62.6
IFNγ 8.75
IFNγ 10.5
IFNγ 11.1
IFNγ 13.3
Comp-PE-A :: IFNg
Comp-BV421-A :: CD3

#### Figure S8

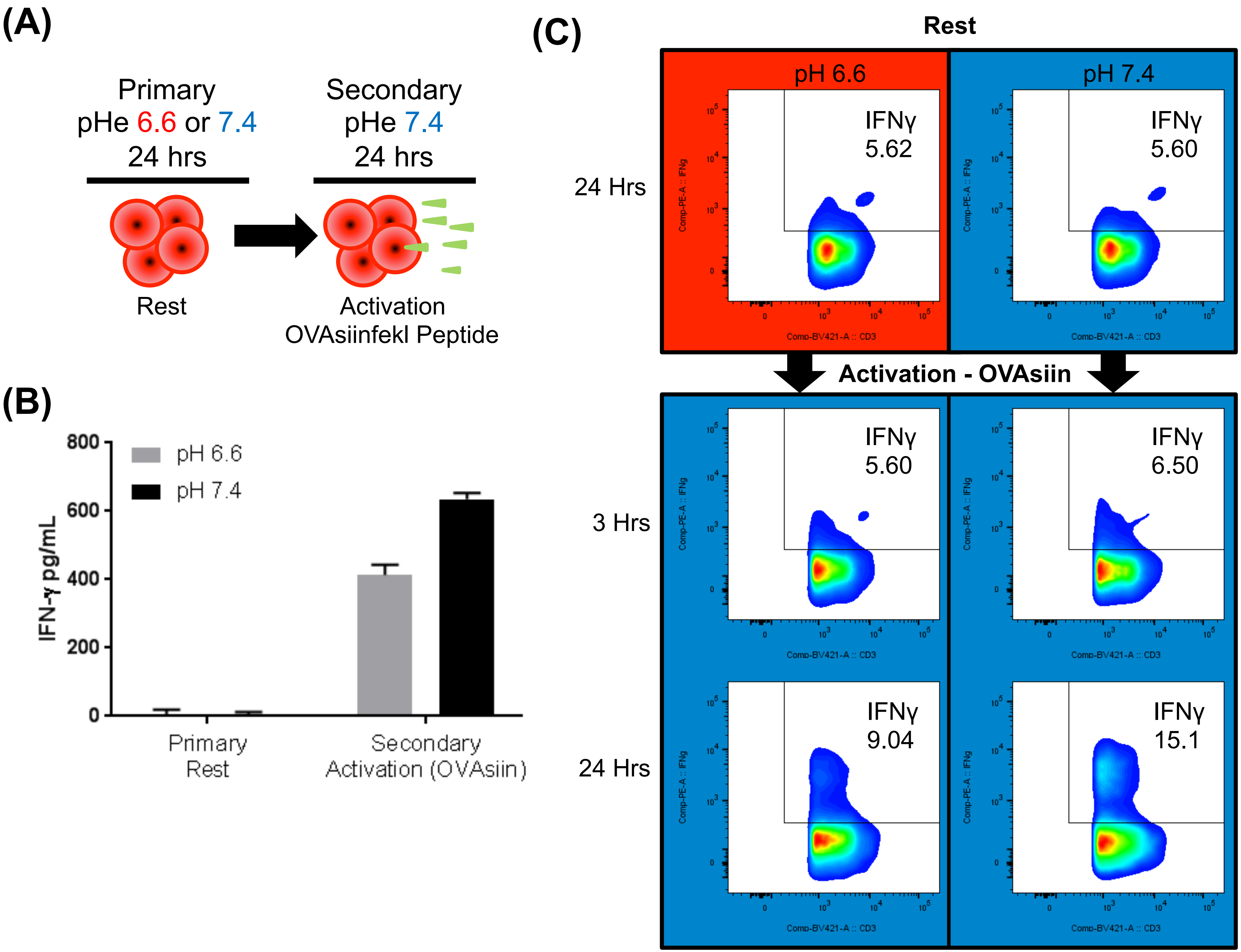
(A)
Primary
pHe 6.6 or 7.4
24 hrs
Rest
Secondary
pHe 7.4
24 hrs
Activation
OVAsiinfekl Peptide
(B)
IFN-γ pg/mL
800
600
400
200
0
pH 6.6
pH 7.4
Primary
Rest
Secondary
Activation (OVAsiin)
(C)
Rest
pH 6.6
pH 7.4
24 Hrs
IFNγ
5.62
IFNγ
5.60
Activation - OVAsiin
3 Hrs
IFNγ
5.60
IFNγ
6.50
24 Hrs
IFNγ
9.04
IFNγ
15.1
Comp-PE-A :: IFNg
Comp-BV421-A :: CD3
